# Microbial dormancy improves predictability of soil respiration at the seasonal time scale

**DOI:** 10.1101/434654

**Authors:** A. Salazar, J.T. Lennon, J.S Dukes

**Author notes:** Corresponding author: Alejandro Salazar.

## Abstract

Climate change is accelerating global soil respiration, which could in turn accelerate climate change. The biological mechanisms through which soil carbon (C) responds to climate are not well understood, limiting our ability to predict future global soil respiration rates. As part of a climate manipulation experiment, we tested whether differences in soil heterotrophic respiration driven by season or climate treatment (R_H_) are linked to 1) relative abundances of microbes in active and dormant metabolic states, 2) net changes in microbial biomass and/or 3) changes in the relative abundances of microbial groups with different C-use strategies. We used a flow-cytometric single-cell metabolic assay to quantify the abundance of active and dormant microbes, and the phospholipid fatty acid (PLFA) method to determine microbial biomass and ratios of fungi:bacteria and Gram-positive:Gram-negative bacteria. R_H_ did not respond to climate treatments but was greater in the warm and dry summer than in the cool and less-dry fall. These dynamics were better explained when microbial data were taken into account compared to when only physical data (temperature and moisture) were used. Overall, our results suggest that R_H_ responses to temperature are stronger when soil contains more active microbes, and that seasonal patterns of RH can be better explained by shifts in microbial activity than by shifts in the relative abundances of fungi and Gram-positive and Gram-negative bacteria. These findings contribute to our understanding of how and under which conditions microbes influence soil C responses to climate.

## Introduction

Every year, microbes from terrestrial ecosystems emit approximately 54 Pg C into the atmosphere via soil heterotrophic respiration (Hashimoto et al., 2015). This is more than five times the amount of C released by fossil fuel emissions in 2016 (ca. 10 Pg C; Le Quéré et al., 2016). Even small increases in this soil C flux, if sustained, could contribute to accelerated global warming. Unless suppressed by dry conditions (Allison and Treseder, 2008; Schindlbacher et al., 2012, Suseela et al. 2012), warming generally increases soil respiration (Carey et al., 2016). Evidence from the last 50 years (i.e., since the first soil respiration records) suggests that soil respiration, both total (i.e., plant roots and soil microbes, R_S_; Bond-Lamberty and Thomson, 2010; Zhao et al., 2017; Bond-Lamberty et al., 2018) and heterotrophic (i.e., microbes in root-free soil, RH; Hashimoto et al., 2015; Bond-Lamberty et al., 2018) have been increasing with temperature at a rate of approximately 0.04 Pg C y^−1^(Zhao et al., 2017); 0.1 Pg C y^−1^ in the last 3 decades (Bond-Lamberty and Thomson, 2010). If changes in global temperature and precipitation regimes accelerate turnover of soil C without proportionally increasing C inputs from plant growth, this would generate a positive soil C-climate feedback.

The mechanisms through which global soil respiration is increasing are not clear, limiting our ability to predict whether this trend is going to accelerate, stabilize, or decrease. Current projections of feedbacks between terrestrial C and climate remain highly uncertain (Friedlingstein et al., 2014), which limits their usefulness to inform climate policy. Recent studies have raised the question of whether or not these uncertainties could be reduced by a better understanding of how microorganisms respond to climatic changes (Bardgett et al., 2008; Wieder et al., 2015).

Two of the best-studied microbial parameters that can be linked to R_H_ are microbial biomass (lleris et al., 2003; Wang et al., 2003; Lee and Jose, 2003; Liu et al., 2009; Zhou et al., 2011) and community composition (Zogg et al., 1997; Monson et al., 2006; Waldrop and Firestone, 2006; Cleveland et al., 2007; Zhou et al., 2011; Don et al., 2017). Assuming that all microbes are active and respiring, one would expect soil R_H_ to increase with the abundance of soil microorganisms. This idea is supported by observations of simultaneous increases or decreases of microbial biomass and R_H_ in response to warming (Zhou et al., 2011), moisture (Illeris et al., 2003; Liu et al., 2009), substrate availability (Wang et al., 2003) and plant biomass removal (Zhang et al., 2005). However, other studies have failed to find a consistent relationship between microbial biomass and R_H_ (Waldrop and Firestone, 2006; Waring and Hawkes, 2015; Birge et al., 2015; Buchkowski et al., 2015), suggesting that microbial parameters besides biomass may contribute to changes in R_H_. Hypotheses linking R_H_ and community composition postulate that a given set of external factors will distinctly favor the growth of particular microbial groups, based on their C needs and C use efficiencies. This idea is supported by observations of, for example, warming increasing the abundance of microbial functional populations specialized in the degradation of labile C, but not recalcitrant C (Zhou et al., 2011).

After microbial biomass and community composition, a third microbial parameter that has been increasingly proposed as an explanatory factor of R_H_ is microbial dormancy (Placella et al., 2012; Manzoni et al., 2014; Wang et al., 2014; Wang et al., 2015; Barnard et al., 2015; He et al., 2015; Salazar et al., 2018). In addition to growing, dying, and changing composition, microbial communities in soil can switch between active and dormant metabolic states (Stenström et al., 2001; Schimel et al., 2007; Lennon and Jones, 2011), for example, in response to warming and drying/wetting cycles (Barnard et al., 2015; Salazar et al., 2018). Changes of metabolic state are generally faster than growth, death, and changes in composition (Blagodatskaya and Kuzyakov, 2013). In part because of this, most experiments exploring the relationship between microbial dormancy and R_H_ have examined short temporal scales (Placella et al., 2012; Aanderud et al., 2015; Barnard et al., 2015; Salazar et al., 2016). Much less in known about the importance of microbial dormancy for R_H_ at longer (e.g., seasonal) temporal scales, which are more relevant for modeling purposes and society-level decision making.

In this study, we investigated whether seasonal R_H_ in a temperate old-field ecosystem is linked to changes in the metabolic state of microbial communities in soil; to net changes in microbial biomass; and/or to changes in the relative abundances of microbial groups that consume and emit soil C at different rates, namely, fungi and bacteria (Six et al., 2006; Sinsabaugh et al., 2016), and Gram-positive and Gram-negative bacteria (Lennon et al., 2012). In addition to the potential changes in total microbial biomass (Devi and Yadava, 2006) and community composition (Waldrop and Firestone, 2006) that can occur on a seasonal time scale, we expected seasonal changes in temperature and moisture to affect the metabolic state of microbial communities in soil. Specifically, we expected the abundance of active microbial biomass to be highest when environmental conditions are optimum for microbial processes, and for this to help explain seasonal changes in soil respiration rates.

## Methods

### Study site

The study was conducted at the Boston-Area Climate Experiment (BACE), located at the University of Massachusetts’ former Suburban Experiment Station in Waltham, Massachusetts (42°23.1’N, 71°12.9’W). The mean annual temperature and precipitation at the site are 10.3 °C and 1063 mm, respectively. The soil at BACE is classified as Mesic Typic Dystrudept (Haven series), with loamy topsoil (45% sand, 46% silt, 9% clay; gravel content: 7%) and a gravelly sandy loam subsoil. The plant community is dominated by non-native grasses and forbs (Hoeppner and Dukes, 2012).

To guarantee that the measured R_H_ was caused by microbial activity and not by plant roots, we collected all of our samples from patches of soil that were isolated from roots and plant carbon inputs by “root-exclusion collars.” These collars were made of 30-cm diameter plastic pipe that had been driven 30 cm into the soil in November 2007 (Suseela et al., 2012). The collars extended ~4 cm above the soil surface. To prevent plant growth within these root-exclusion collars, we covered the soil surface within each collar with a circle of weed-blocking nylon mesh. This mesh was removed only during R_H_ measurements and soil sampling. Carbon inputs had been limited in this manner for the previous nine years; by the fourth year of plant exclusion (2011), labile organic matter remaining in the soils was already substantially depleted in comparison to the surrounding soils in which plants grew, as shown by lower rates of substrate-induced respiration (Koyama et al. 2018). Thus, our use of these root-free soils with similar past C inputs enabled a controlled examination of microbial responses, but the sustained lack of plant inputs and the consequently depleted labile organic matter need to be kept in mind when interpreting our results.

### Experimental design

The BACE manipulated climatic conditions in 36 square experimental plots, each 2 m × 2 m. A factorial combination of four levels of warming and three levels of precipitation created a total of 12 climate treatments. The experiment consisted of three replicate blocks. Within each block, plots were arranged linearly in three groups of four, with each group receiving one of the three precipitation treatments. The four plots within each group were spaced 1 m apart, with one plot receiving each of the four levels of warming. Each block was located under a single greenhouse frame that served as a mount for infrastructure related to the precipitation treatments.

Warming was applied with ceramic infrared heaters mounted 1 m above each corner of each plot, and facing towards the center of the plot and down at a 45° angle. The treatments corresponded to the wattage of the heaters surrounding each plot: unheated (0 W), low (200 W), medium (600 W), and high (1000 W) heat. The three heated plots within each group were wired to a single circuit, and the warming system was programmed to attempt to maintain a 4 °C difference between the canopy temperatures of the unheated and high heat plots within each group. The power supplied to the heaters in each group was adjusted every 10 s based on the measured temperature difference between the unheated and high heat plots in that group. Canopy temperatures were measured with infrared radiometers (IRR-PN; Apogee Instruments, Logan,UT, USA) placed at a 45° downward angle, 1 m above the northern edges of the plots. The four warming levels approximately simulated the different warming scenarios projected by the Intergovernmental Panel on Climate Change (IPCC) for the end of this century (Stocker, 2014).

The precipitation manipulation included ambient, dry (−50% of all precipitation year-round), and wet (+50% growing season rainfall) treatments. Above the dry treatment, rainfall was captured by 15 cm-wide clear polycarbonate slats spaced 15 cm apart that were mounted on the greenhouse frames, >2 m off the ground. From early May to mid-November, the removed rainfall was collected in tanks and immediately applied to the wet treatments with a sprinkler system. Further details of the experiment can be found in Hoeppner and Dukes (2012), Suseela et al., (2012), and Auyeung et al., (2013).

### *In situ* soil measurements and sampling

We made *in situ* soil measurements (R_H_, and microbial activity/dormancy) and collected samples for analysis in the laboratory in the summer (June) and fall (October) of 2016. For simplicity, we refer to these measurements by the season in which they were made, but it is important to recognize that they represent snapshots of the conditions at the moment of sampling and not an average of the respective months or seasons.

We measured R_H_ with a LICOR 6400 soil CO2 flux chamber (LI-COR Biosciences, Inc. Lincoln, NE, USA), in small PVC collars (10 cm in diameter and 5 cm in height, 2 cm into the soil) that we had installed within the root-exclusion collars in April 2016. During the R_H_ measurements we also measured soil temperature (10 cm) using a thermocouple probe. We measured volumetric soil moisture on the same day using time-domain reflectometry waveguides installed vertically (0-10 cm depth) in the root-exclusion collars.

We used a soil sampler (2 cm diameter) to collect soil from the top 10 cm immediately after measuring R_H_. We used 1 g of the soil for measuring *in situ* active and dormant microbial biomass (see below). We stored the rest of the soil in coolers with ice packs and transported them to Purdue University where we measured microbial biomass and fungi:bacteria and Gram-positive-Gram-negative ratios (see below). At Purdue, samples were stored at 4 °C until all samples were processed.

### Abundance of active and dormant microbes in soil

We used a flow-cytometric single-cell metabolic assay to quantify active and dormant microbial abundance (del Giorgio and Gasol, 2008). Immediately after measuring R_H_ and collecting soil samples in each plot, we mixed 1 g of soil with 9 mL of distilled, sterile water and vortexed this solution for 1 min. We filtered the solution (particle retention > 11 μm) to remove large debris. We sampled 0.8 mL of the filtered solution and added 0.1 mL of 4’,6-diamidino-2-phenylindole (DAPI; 5 μg DAPI mL^−1^ final concentration) and 0.1 mL of 5-cyano-2,3-ditolyl tetrazolium chloride (CTC; 5 mM final concentration) to stain all and respiring-only cells, respectively. DAPI stains the DNA of all viable cells while only metabolically active cells can transform the electron acceptor CTC to the fluorophore CTC-formazan (Kaprelyants and Kell, 1993). We mixed this solution in a shaker for 30 min and then stopped the reaction by storing the samples in coolers with ice packs. Immediately after processing all samples, we shipped them to the flow cytometry facility at Indiana University, USA. To check for potential auto-fluorescence in soil, we also analyzed negative-control unstained samples for each plot.

To estimate the abundance of active (i.e., DAPI and CTC co-labeled) and dormant (i.e., DAPI-only labeled) cells, we used an LSRII flow cytometer (Becton-Dickinson, San Jose, CA) equipped with Forward Scatter PMT (FSC-PMT) for improved resolution of small particles. We used the FACSDiva v.6.1.3 software for data analysis. DAPI was excited with a 20mW 405nm laser, and detected using a 450/50 filter, while CTC was excited with a 30mW 488nm laser, and detected using a 695/40 filter. To further resolve small particles, we set the LSRII window extension at 2.00 (rather than the default of 10.00). To minimize the amount of debris and background included in the sample analysis, we set thresholds at 1000 and 750 for FSC-PMT and SSC parameters, respectively. We ran controls (unlabeled, DAPI, and CTC) for each sample, and saved 10,000 events per sample. The sample injection was rinsed after each sample in order to minimize any cross contamination between samples. An example of the analysis output (Figure S1) is shown in the supplementary material.

We acknowledge three caveats with our approach for measuring abundance of active and dormant cells: 1) we did not measure dead cells, so our estimate of dormant cells could be overestimated by non-CTC stained cells that were not viable, 2) a fraction of the dormant cells in soil could have been activated when suspending samples in water for dye application, which would underestimate the proportion of inactive cells, and 3) Because CTC stains bacteria but not fungi and because we measured abundance of CTC/DAPI labeled cells based on light scattering characteristics of *Escherichia coli* (see supplementary material), our measurements of microbial activity reflect bacteria but not fungi. Based on these caveats, we made the following assumptions: We assumed that the cell structure of most dead cells was compromised, preventing the cells from retaining DAPI-labeled DNA and/or affecting its light scattering characteristics, and we therefore assumed that most dead cells were not counted as dormant. We also assumed that any activation of dormant cells during the exposure to the dyes was minimal (which seems reasonable considering the low fractions of active bacteria – see Results) and similar for all samples (i.e., allowing comparisons among treatments). Because bacteria dominated the root-free soils considered in this study (see Results), we suspect that measurements of bacterial activity are a reasonable indicator of overall soil microbial activity.

### Microbial biomass and relative abundances of microbial groups

We measured microbial biomass, fungi:bacteria ratios, and Gram-positive:Gram-negative ratios with the phospholipid fatty acid (PLFA) method (Hurst et al., 1997). We estimated microbial biomass based on analysis of phospholipid phosphates (PLPO4) and fungi:bacteria and Gram-positive:Gram-negative ratios based on analysis of phospholipids fatty acids (as in Acosta-Martinez et al., 1999). We extracted lipids from 5 g of soil (bags stored at 4 °C) using a chloroform/methanol/phosphate buffer and fractionated phospholipids using column chromatography. We measured PLPO4 colorimetrically at 610 nm (DU^®^730 UV/VIS spectrophotometer, Beckman Coulter, Inc., Fullerton, CA) and fatty acids via Gas Chromatography-Mass Spectrometric Detection (Agilent 7890, Agilent 5975 MSD, Agilent Technologies Inc., Santa Clara, CA).

We calculated fungi:bacteria ratio as the predominant fungal PLFAs 18:2w6, divided by the sum of the predominant bacteria PLFAs 14:0, i15:0, a15:0, 15:0, 16:0, 10Me16:0, i17:0, a17:0, cy17:0, Me18:0, and cy19:0 (Bååth and Anderson, 2003). Similarly, we calculated Gram-positive:Gram-negative ratios as the sum of Gram-positive PLFAs i15:0, a15:0, i17:0, and a17:0, divided by the sum of Gram-negative PLFAs cy19:0 and cy17:0 (Joynt et al, 2006).

The PLFA method does not allow a highly resolved analysis of the composition and structure of microbial communities in soil. However, previous studies have shown that respiration responses to the environments are conserved at fairly coarse phylogenetic scales (Lennon et al., 2012).

### Statistical analysis

We first analyzed the effects of the warming and precipitation treatments (fixed effects) on R_H_, using a mixed-effects model that included time as a fixed effect and block as a random effect. For this we used the *lmer* function from the *lme4* package (Bates et al., 2014) in R, version 3.3.1. We then used a multiple correlation analysis (*lm* function) to estimate how much of the seasonal (fixed effect) differences in R_H_ were explained by soil temperature, moisture, Total Microbial Biomass (TMB), Active Microbial Biomass (AMB), and the relative abundance of microbial groups (i.e., fungi:bacteria and Gram-positive:Gram-negative ratios). We used the *glmulti* function, from the *glmulti* package (Calcagno and de Mazancourt, 2010), to select the best statistical model. We compared models based on the Bayesian Information Criterion (BIC), which accounts for differences in the number of explanatory variables among models.

## Results

The warming and precipitation treatments affected environmental conditions, but had little effect on R_H_ in these relatively dry soils. Instead, R_H_ differed across seasons. These seasonal differences in R_H_ were explained better by temperature and the abundance of actively respiring cells than by environmental or microbial variables alone.

### Effect of experimental warming on soil temperature

The warming treatments increased (P<0.05; Table S1) soil temperature in both seasons (Figure 1 a and b). Soil temperature was affected by the precipitation treatments as well. Soil temperature was higher (P<0.05) in the dry (and less plant-shaded) plots than in the ambient and wet plots, especially in the Fall (P = 0.06). In the summer, soil temperature ranged from 21.5 ± 1.2 °C in the unheated plots to 23.9 ± 2.0 °C in the high heated plots. In the fall, soil temperature ranged from 15.4 ± 1.5 °C in the unheated plots to 17.9 ± 1.7 °C in the high heated plots. Differences in soil temperature between unheated and high heated plots were ca. 2.5 °C in both seasons, while differences between seasons were in average ca. 6 °C. Overall, differences in soil temperature were larger between seasons than across warming treatments.

**Figure 1.**
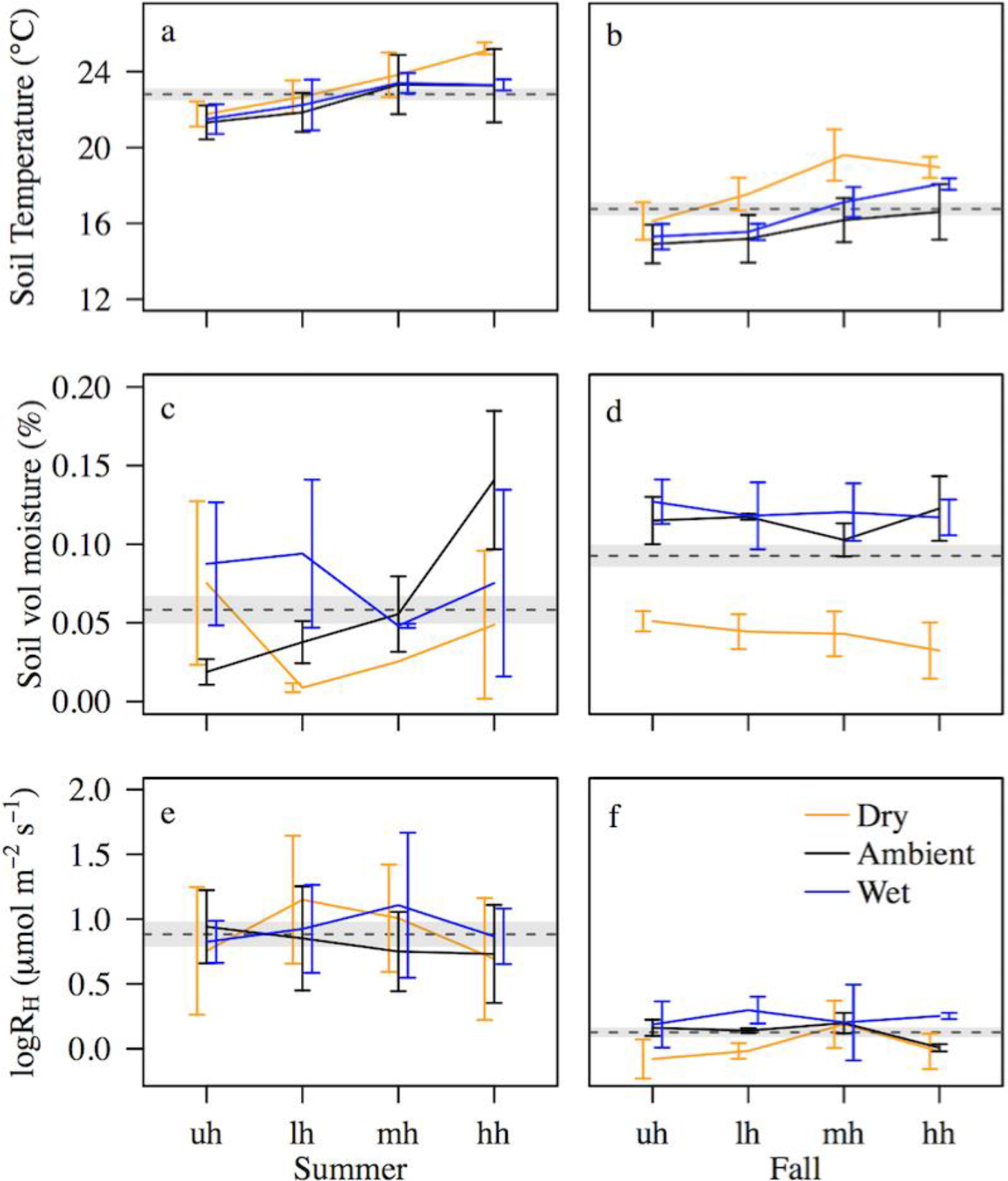
Soil temperature (a,b), moisture (c,d), and R_H_ (e,f) in summer (a,c,e) and fall (b,d,f) across warming and precipitation treatments. uh: unheated, lh: low heat, mh: medium heat, and hh: high heat. Statistics in Tables S1-3. Values are means ± SE. Dashed lines and shaded areas indicate averaged ± SE values in each season. No error bar is shown in the mh-dry treatment in panel *c* because of missing data.

### Effect of precipitation manipulation on soil moisture

Although there were differences (P < 0.05) in soil moisture across seasons and precipitation treatments (Figure 1 c and d; Table S2), soils were fairly dry (<20% v/v) in all cases. Differences in soil moisture across treatments were larger (P = 0.02) in the fall than in the summer. In the fall, soils from the dry treatment (ca. 5% v/v) were drier than those from the ambient and wet plots (ca. 12% v/v in both precipitation treatments across all warming treatments). Averaged across treatments, soils were drier in the summer (6% v/v) than the fall (9% v/v; P<0.01) (see also Figure 2c).

**Figure 2.**
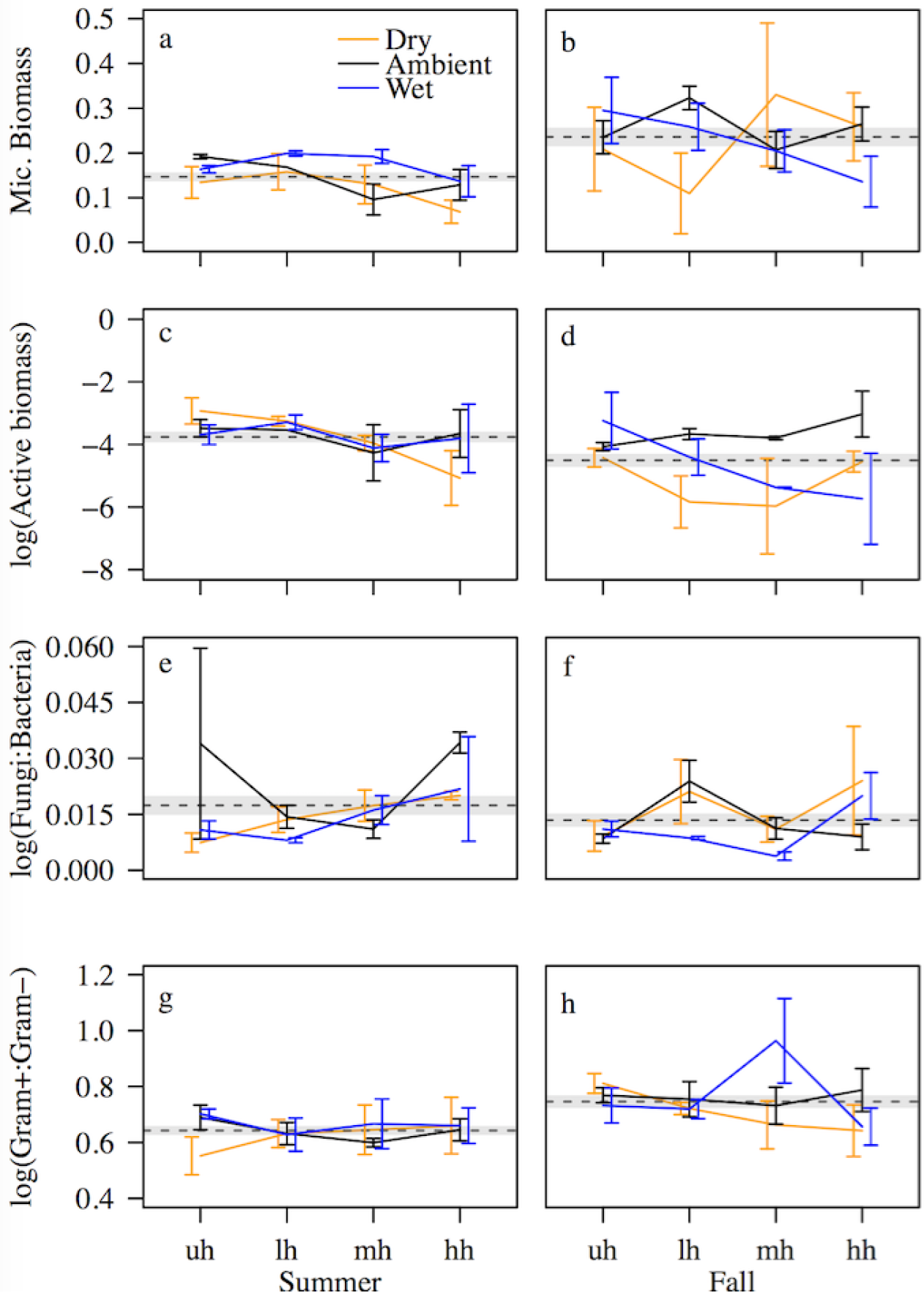
Microbial biomass (a,b; in phospholipid phosphate g^−1^soil), Active biomass (c,d; in phospholipid phosphate g^−1^soil), Fungi:Bacteria ratios (e,f), and Gram positive:Gram negative ratios (g,h), in summer (a,c,e,g) and fall (b,d,f,h) across warming and precipitation treatments. uh: unheated, lh: low heat, mh: medium heat, and hh: high heat. Statistics in Tables S4-7. AMB, and Fungi:Bacteria and Gram positive:Gram negative ratios in statistical models were log transformed to meet assumptions. Values are means ± SE. Dashed lines and shaded areas indicate averaged ± SE values in each season. No error bars in lh-ambient treatment in panels *a* and *c* because of missing data.

### Effects of warming and precipitation treatments on R_H_

R_H_ did not differ (P > 0.05) across the warming and precipitation treatments but it differed (P<0.05) between seasons (Figure 1 e and f; Table S3). In the summer, R_H_ averaged 2.88 ± 0.30 μmol m^−2^s^−1^across all the warming and precipitation treatments. By the fall, average R_H_ (1.16 ± 0.05 μmol m^−2^s^−1^) had decreased (P<0.05) by more than 50%. Although in the fall R_H_ tended to increase from dry to wet plots (Figure 1f), this trend (like all other differences across treatments) was not statistically significant (Table S3).

### Effects of warming and precipitation treatments on microbial parameters

Microbial parameters differed between seasons but were unaffected or weakly affected by treatments (Figure 2 and Tables S4-7). Although microbial biomass responded to warming under some precipitation treatments (i.e., marginal interaction, P = 0.07; Figure 2 a and b; Table S4), and active microbial biomass and fungi:bacteria ratios marginally (P = 0.05 and P = 0.06, respectively) increased with warming, differences in microbial parameters across treatments were small in comparison with the marked differences between seasons. From summer to fall, microbial biomass and Gram positive:Gram negative ratios increased (P < 0.05) by 60% (Figures 2 a and b; Table S4) and 16% (Figures 2 g and h; Table S7), respectively; fungi:bacteria ratios did not change (Figure 2 e and f; Table S6); and active microbial biomass decreased (P < 0.05) by 40% (Figure 2 c and d; Table S5).

### Predictors of seasonal R_H_

Seasonal differences in R_H_ were primarily explained by temperature and the abundance of metabolically active microbes in soil (Table 1). The statistical model that best fitted our data (P < 0.05, BIC = 69.7, Table S4) surprisingly suggests that decreases in R_H_ between the summer and fall were associated with increases (P < 0.05) in TMB, TMB having a different influence on R_H_ in each season (P < 0.05). For reasons that we discuss below, we also analyzed the second best statistical model (P < 0.05, BIC = 81.5), which suggests that seasonal decreases in R_H_ from summer to fall were associated with decreases in temperature and in the abundance of actively respiring cells in soil (Table 1).

**Table 1.** Statistics of the best explanatory model for seasonal R_H_. Significance codes: P < 0.001 ‘***’, 0.001 < P < 0.01 ‘**’.

|  | **Estimate** | **Std. Error** | **t-value** | **P** |
| --- | --- | --- | --- | --- |
| Intercept | -0.894 | 0.614 | -1.455 | 0.152 |
| Temp | 0.107 | 0.025 | 4.254 | < 0.001*** |
| Temp:Moisture | -7.429 | 6.058 | -1.226 | 0.226 |
| Temp:log(AMB) | 0.008 | 0.003 | 2.730 | 0.009** |

Overall, temperature and moisture explained seasonal soil respiration better than microbial processes alone, but incorporation of microbial data increased explanatory power (Figure 3). Our results suggest that, on average, log(R_H_) increased by 0.11 μmol m^−2^s^−1^per 1 °C increase in soil temperature (Table 1) and that the magnitude of this response increased with the abundance of active microbes in soil (Figure 4). This model explained 35% (adjusted R^2^) of the variation in R_H_.

**Figure 3.**
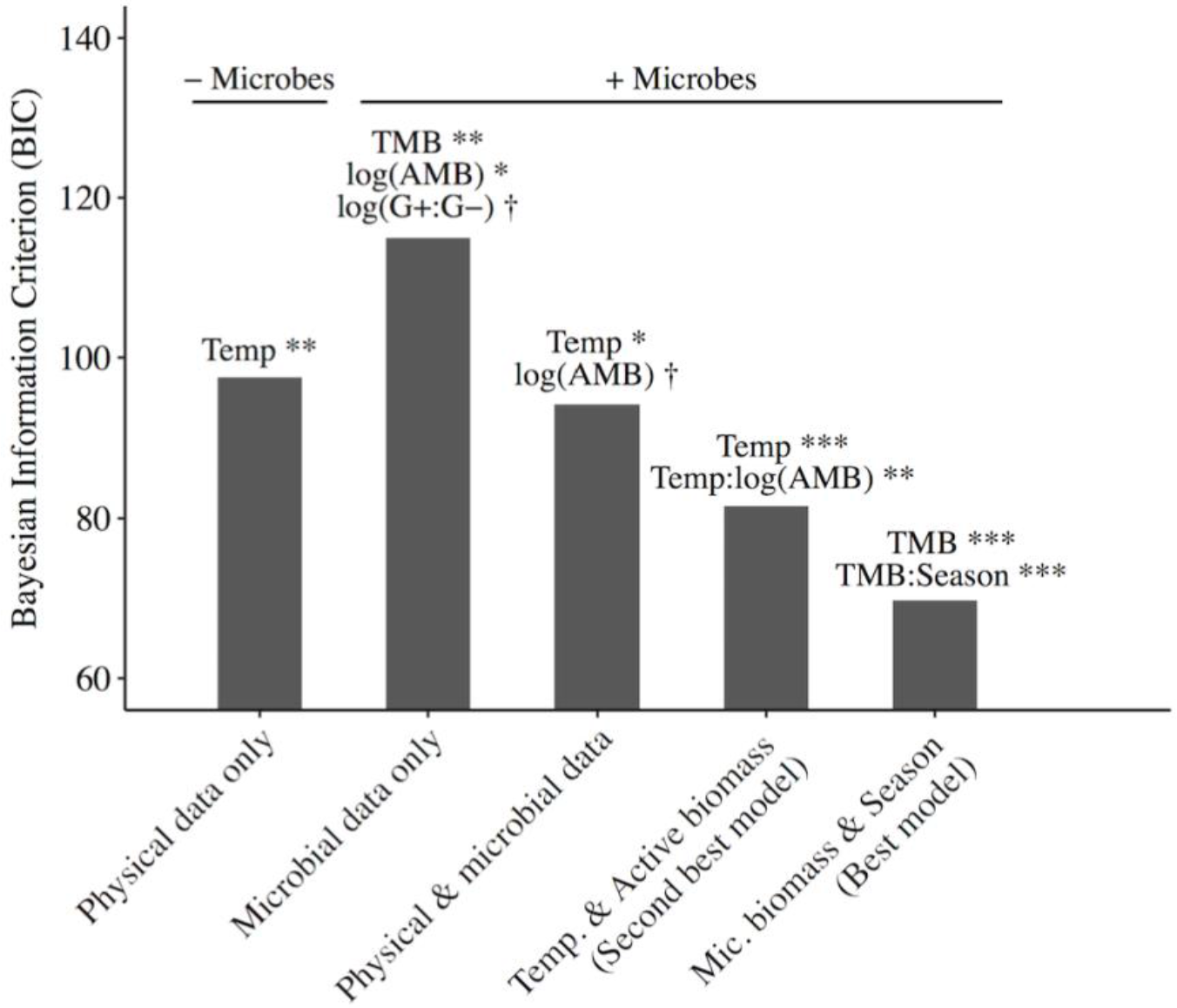
Goodness of fit, as measured by the Bayesian Information Criterion (BIC), of models of log(R_H_) that include different categories of explanatory variables. Lower scores indicate better model fits. Categories include models fitting log(R_H_) as a function of only physical conditions (as a function of temperature and moisture; statistics in Table S10); only values related to microbes: TMB, log(AMB), log(Fungi:Bacteria), and log(Gram-positive:Gram-negative); statistics in Table S9); with microbes and physical conditions (as a function of temperature, moisture, TMB, log(AMB), log(Fungi:Bacteria), and log(Gram-positive:Gram-negative); statistics in Table S11); the second best model (as a function of temperature and the interaction between temperature and log(AMB); statistics in Table 1); and the best (but see discussion) model (as a function of TMB, and the interactions between log(AMB) and moisture, and TMB and season; statistics in Table S8). Significance codes: *P* < 0.001 ‘***’, 0.001 < *P* < 0.01 ‘**’, 0.01 < *P* < 0.05 ‘*’, 0.5 < *P* < 0.1 ‘†’.

**Figure 4.**
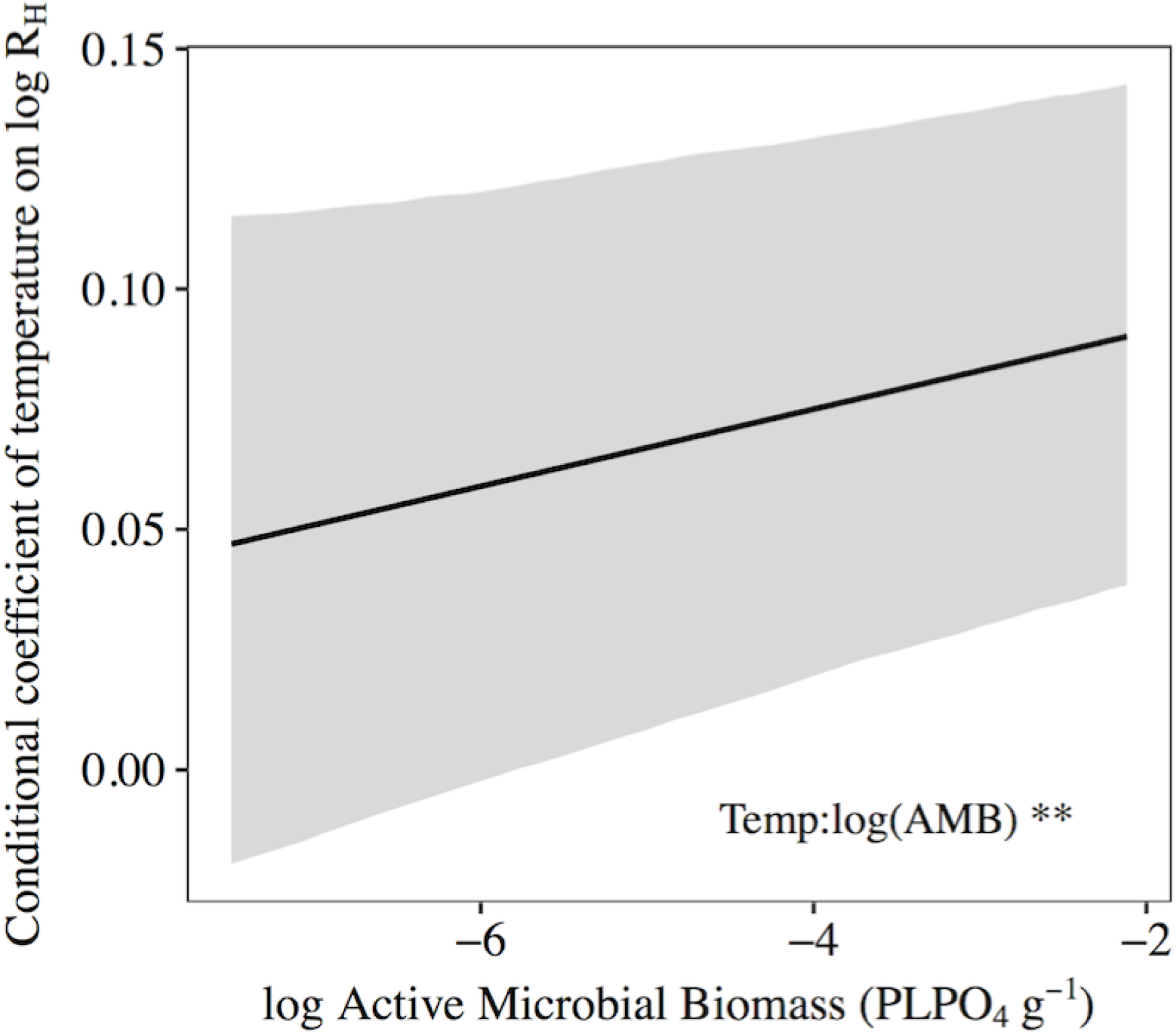
Changes in the coefficient of soil temperature, in the two-way interaction term with AMB (Table 1), conditional on AMB (*interplot* function in R). AMB in statistical model was log transformed to meet assumptions. Grey area indicates 95% confidence intervals.

## Discussion

Recently, the policy relevance of soil C-climate feedbacks has motivated a wide range of research on mechanisms that regulate soil C cycling, and their associated temporal scales. One of those mechanisms is the metabolic activation and deactivation of soil microbes in response to favorable and stressful environmental conditions. Previous studies have demonstrated that soil respiratory responses to temperature and moisture at a temporal scale of hours to days are associated with microbes switching between active and dormant metabolic states (Placella et al., 2012; Barnard et al., 2015; Salazar et al., 2016). However, less is known about the importance of these mechanisms over longer timescales. The results of this study suggest that the abundance of active microbes in soil changes at the seasonal scale too, and that these shifts affect soil respiration rates.

The abundances of total and active microbes in soil can change across seasons. In our study, AMB (and R_H_) was greater in the summer than in the fall. However, TMB was greater in the fall than in the summer. Increases in TMB between June and October could have been partially caused by increases in soil moisture, which likely facilitated access of microbes to nutrients. Increases in soil moisture during the still warm end of the summer could have stimulated microbial growth, but decreases in temperature in the fall likely induced a large proportion of microbes in the soil to enter dormancy. This could explain why AMB was higher in the summer than in the fall, even though TMB was lower. We know of only one other study that simultaneously monitored TMB and AMB at the seasonal scale. In it, Van de Werf and Verstraete (1987) found TMB to increase by 29% from June to August in a fallow topsoil, while AMB remained practically unchanged (Van de Werf and Verstraete, 1987). However, in winter-wheat soil both TMB and AMB increased from June to August (Van de Werf and Verstraete, 1987). This suggests that seasonal changes in TMB and AMB can also be affected by soil type and/or agricultural practices (see also Girvan et al., 2003). Similarly, in the first year of a two-year warming study in a temperate forest (Schindlbacher et al., 2011), microbial biomass increased (both in heated and unheated treatments) by 30% from July to September, while microbial metabolic activity (measured as soil respiration rates per concentration of microbial biomass) decreased by 50%. Interestingly, in the second year, these trends reversed (Schindlbacher et al., 2011). Together, our results and these observations show that the amount of total and active microbial biomass in soil do not necessarily change in the same direction and at the same time across seasons.

Seasonal changes in microbial biomass can happen in parallel with changes in community composition. In our study, increases in TMB (and decreases in AMB) between summer and fall were accompanied, on average, by a decrease in fungi:bacteria ratios (Figure 2 e and f) and an increase in Gram positive:Gram negative ratios (Figure 2 g and h). The change in fungi:bacteria ratio could have been caused by warmer summer temperatures favoring fungi over bacteria (Zhang et al., 2005, Castro et al., 2010) and/or by faster bacterial growth between the summer and the fall as soil moisture increased (Figure 1 c and d). Although fungi play a key role in soil C cycling in some systems such as nutrient-rich forest soil (Baldrian et al., 2012), the root-free soil from this experiment was dominated by bacteria (fungi:bacteria ratios were always < 0.1). This suggests that changes within the bacterial community may have been more important for soil C cycling than relative changes in the abundance of fungi and bacteria.

Our multiple regression analysis suggests two alternative, and possibly complementary, explanations for why the microbial parameters discussed above contribute to seasonal R_H_. The statistical model that best fitted our data suggests a relationship between seasonal decreases in R_H_ with increases in TMB. An inverse relationship between TMB and R_H_ could reflect pulses of R_H_ caused by active microbes recycling necromass C (Geyer et al., 2016). It is plausible that the more severe dryness in the summer than in the fall in our study, led to elevated microbial mortality and therefore to a larger abundance of necromass C accessible to active microbes. However, given the capacity of microbes to adjust their metabolism and remain viable under stressful conditions, the contributions of cell lysis to soil C fluxes is probably insignificant (Halverson et al., 2000). We do not know of any other biological process that could explain this result and therefore recommend caution when interpreting its causality. On the other hand, the statistical model that provided the second-best fit to our data suggests that seasonal decreases in R_H_ from summer to fall were driven by decreases in soil temperature and in the abundance of metabolically active microbes in soil. This is consistent with theory of microbial physiology (Stenström et al., 2001; Schimel et al., 2007; Lennon and Jones, 2011) and with experiments conducted at short temporal scales (Placella et al., 2012; Aanderud et al., 2015; Barnard et al., 2015; Salazar et al., 2016). If our one-time measurements of active/dormant biomass from summer and fall are close to the respective seasonal averages, our results would indicate that temperature and microbial metabolism data alone are powerful in predicting seasonal R_H_.

Although Gram-positive:Gram-negative ratios did not contribute to the best-fitting models of R_H_ (Tables 1 and S8), changes within the bacterial community could help to explain the relationship between AMB and R_H_. Increases in Gram-positive:Gram-negative ratio from summer to fall could have been associated with different capabilities of the bacterial groups to cope with moisture stress. Gram-positive bacteria have a peptidoglycan-rich cell wall that makes them more resistant to dry conditions than Gram-negative bacteria (Halverson et al., 2000; Fuchslueger et al., 2014). Considering that soils in our experiment were relatively dry (< 20% v/v) in both seasons, it is likely that Gram-negative bacteria were more severely affected by moisture stress than their thick-cell-wall counterparts. Some Gram-negative bacteria have higher maximum respiration rates than Gram-positive bacteria (e.g. Acidobacteria vs. Actinobacteria, respectively; Lennon et al., 2012). It is possible that from summer to fall there were larger decreases in the abundance of metabolically active microbes with high maximum respiration rates but low resistance to dryness (e.g. Acidobacteria), relative to bacterial groups with low maximum respiration rates but high resistance to dryness (e.g. Actinobacteria). However, not all Gram-negative bacteria have higher maximum respiration rates than Gram-positive bacteria. Gram-positive Firmicutes have higher maximum respiration rates than Gram-negative Acidobacteria, Bacteroidetes and Proteobacteria (Lennon et al., 2012). Therefore, decreases in R_H_ between summer and fall could also have been associated with metabolic deactivation of microbes with high maximum respiration rates and high resistance to dryness. We would need a more resolved composition analysis to know which (if any) of these alternative explanations was the case in our study. Nonetheless, our results suggest that, as soil moisture levels change, the abundance of microbial groups with different levels of resistance to dryness could influence the size of the microbial pool that remains metabolically active.

Finally, our findings suggest that the effect of temperature on R_H_ gets stronger with the abundance of metabolically active microbes in soil. Microbes that are pushed to enter dormancy by moisture stress are practically unaffected by changes in temperature. However, the metabolic rates (e.g. respiration) of active microbes are sensitive to temperature (Anderson and Domsch, 1985) and therefore it is reasonable to expect that a greater abundance of active microbes in soil makes R_H_ more sensitive to temperature. This builds on previous observations of R_H_ being less sensitive to warming (and precipitation) under dry, presumably water-stressed conditions (Schindlbacher et al., 2012, Suseela et al., 2012; Koyama et al. 2018).

In summary, we found that 1) seasonal changes in total microbial biomass in soil do not necessarily reflect changes in the amount of microbial biomass that is metabolically active and capable of driving soil C processes, 2) the metabolic state of soil microbial communities can be more important for seasonal R_H_ than the relative abundances of microbial groups such as fungi and bacteria (Gram-positive and Gram-negative), and 3) the magnitude of the temperature effect on R_H_ increases with the abundance of metabolically active microbes in soil. This work builds on recent research distinguishing active from dormant microbes and highlighting the importance of the metabolically active community for microbe-driven processes. Although few studies to date have linked microbial metabolic state patterns with rates of soil CO2 efflux, our findings suggest the possibility that recent increases in global soil respiration rates could be linked to climate-driven increases in the abundance of metabolically active microbes in soil.

## Acknowledgments

We gratefully acknowledge the help of Clara Vasquez and Risa McNellis with the fieldwork at BACE, the help of Elizabeth Morgan Davis and Nishit Banka with the PLFA analyses, and the help of Christiane Hassel with the flow cytometry analyses. We further acknowledge the many technicians and assistants who constructed and maintained the BACE. AS acknowledges COLCIENCIAS (Departamento Administrativo de Ciencia Tecnología e Innovación en Colombia) and the Fulbright-Colombia program. In addition, this work was supported in part by National Science Foundation Dimensions of Biodiversity Grant 1442246 (JTL) and US Army Research Office Grant W911NF-14-1-0411 (JTL). Work at the BACE was made possible by funding from the United States Department of Agriculture (USDA; 2015-67003-23485; JSD). The BACE was built and maintained with funding from the National Science Foundation (DEB-0546670 and DEB-1146279) and the US Department of Energy’s Office of Science (BER), through the Northeastern Regional Center of the National Institute for Climatic Change Research, and the Terrestrial Ecosystem Science program. We thank the University of Massachusetts and UMass Extension for leasing land to the BACE. Partial support for JSD’s participation in this project was provided by Hatch project 1000026 of the USDA’s National Institute of Food and Agriculture. This is paper no. 1801 of the Purdue Climate Change Research Center (PCCRC). We declare no conflict of interest.

## References

Aanderud Z, Jones S, Fierer N, Lennon JT (2015) Resuscitation of the rare biosphere contributes to pulses of ecosystem activity. Front Microbiol 6: 24. https://doi.org/10.3389/fmicb.2015.00024

Acosta-Martinez V, Reicher Z, Bischoff M, Turco RF (1999) The role of tree leaf mulch and nitrogen fertilizer on turfgrass soil quality. Biol Fertil Soils 29: 55–61. https://doi.org/10.1007/s003740050

Allison SD, Treseder KK (2008) Warming and drying suppress microbial activity and carbon cycling in boreal forest soils. Global Change Biol 14:2898–2909. https://doi.org/10.1111/j.1365-2486.2008.01716.x

Anderson TH, Domsch KH (1985) Determination of ecophysiological maintenance carbon requirements of soil microorganisms in a dormant state. Biol Fertil Soils 1:81–89.

Auyeung DS, Suseela V, Dukes JS (2013) Warming and drought reduce temperature sensitivity of nitrogen transformations. Global Change Biol 19:662–676. https://doi.org/10.1111/gcb.12063

Bååth E, Anderson T-H (2003) Comparison of soil fungal/bacterial ratios in a pH gradient using physiological and PLFA-based techniques. Soil Biol Biochem 35:955–963. https://doi.org/10.1016/S0038-0717(03)00154-8

Baldrian P, Kolařík M, Štursová M, Kopecký J, Valášková V, Větrovský T, Žifčáková L, Šnajdr J, Rídl J, Vlček Č, Voříšková J (2012) Active and total microbial communities in forest soil are largely different and highly stratified during decomposition. ISME J 6:248–258. http://doi.org/10.1038/ismej.2011.95

Bardgett RD, Freeman C, Ostle NJ (2008) Microbial contributions to climate change through carbon cycle feedbacks. ISME J 2:805–814. http://doi.org/10.1038/ismej.2008.58

Barnard RL, Osborne CA, Firestone MK (2015) Changing precipitation pattern alters soil microbial community response to wet-up under a Mediterranean-type climate. ISME J 9:946–957. http://doi.org/10.1038/ismej.2014.192

Bates D, Maechler M, Bolker B, Walker S, Christensen RHB, Singmann H, Dai B, and Grothendieck G (2014) Package ‘lme4’. R foundation for statistical computing, Vienna, 12.

Birge HE, Conant RT, Follett RF, Haddix ML, Morris SJ, Snapp SS, Wallenstein MD, Paul EA (2015) Soil respiration is not limited by reductions in microbial biomass during long-term soil incubations. Soil Biol Biochem 81:304–310. https://doi.org/10.1016/j.soilbio.2014.11.028

Blagodatskaya E, Kuzyakov Y (2013) Active microorganisms in soil: Critical review of estimation criteria and approaches. Soil Biol Biochem 67:192–211. https://doi.org/10.1016/j.soilbio.2013.08.024

Bond-Lamberty B, Thomson A (2010) Temperature-associated increases in the global soil respiration record. Nature 464:579–582. http://doi.org/10.1038/nature08930

Bond-Lamberty B, Bailey VL, Chen M, Gough CM, Vargas R (2018). Globally rising soil heterotrophic respiration over recent decades. Nature 560:80–83. https://doi.org/10.1038/s41586-018-0358-x

Buchkowski RW, Schmitz OJ, Bradford MA (2015) Microbial stoichiometry overrides biomass as a regulator of soil carbon and nitrogen cycling. Ecology 96:1139–1149. https://doi.org/10.1890/14-1327.1

Calcagno V, de Mazancourt C (2010) glmulti: an R package for easy automated model selection with (generalized) linear models. J Stat Softw 34:1–29.

Carey JC, Tang J, Templer PH, Kroeger KD, Crowther TW, Burton, AJ, Dukes JS, Emmett B, Frey SD, Heskel MA, Jiang L (2016) Temperature response of soil respiration largely unaltered with experimental warming. Proc Natl Acad Sci USA 113:13797–13802. https://doi.org/10.1073/pnas.1605365113

Castro HF, Classen AT, Austin EE, Norby RJ, Schadt CW (2010) Soil microbial community responses to multiple experimental climate change drivers. Appl Environ Microbiol 76:999–1007. http://doi.org/10.1128/AEM.02874-09

Cleveland CC, Nemergut DR, Schmidt SK, Townsend AR (2007) Increases in soil respiration following labile carbon additions linked to rapid shifts in soil microbial community composition. Biogeochemistry 82:229–240. http://doi.org/10.1007/s10533-006-9065-z

del Giorgio PA, Gasol JM (2008) Physiological Structure and Single-Cell Activity in Marine Bacterioplankton. In: Kirchman, DL (ed) Microbial Ecology of the Oceans, 2nd edn. Wiley, New Jersey, pp 243–298.

Devi NB, Yadava PS (2006) Seasonal dynamics in soil microbial biomass C, N and P in a mixed-oak forest ecosystem of Manipur, North-east India. Appl Soil Ecol 31:220–227. https://doi.org/10.1016/j.apsoil.2005.05.005

Don A, Böhme IH, Dohrmann AB, Poeplau C, Tebbe CC (2017) Microbial community composition affects soil organic carbon turnover in mineral soils. Biol Fertil Soils 53:445–456. http://doi.org/10.1007/s00374-017-1198-9

Friedlingstein P, Meinshausen M, Arora VK, Jones CD, Anav A, Liddicoat SK, Knutti, R (2014) Uncertainties in CMIP5 climate projections due to carbon cycle feedbacks. J Clim 27:511–526. https://doi.org/10.1175/JCLI-D-12-00579.1

Fuchslueger L, Bahn M, Fritz K, Hasibeder R, Richter A (2014) Experimental drought reduces the transfer of recently fixed plant carbon to soil microbes and alters the bacterial community composition in a mountain meadow. New Phytol 201:916–927. https://doi.org/10.1111/nph.12569

Geyer KM, Kyker-Snowman E, Grandy AS, Frey SD (2016) Microbial carbon use efficiency: accounting for population, community, and ecosystem-scale controls over the fate of metabolized organic matter. Biogeochemistry 127:173–188. https://dx.doi.org/10.1007/s10533-016-0191-y

Girvan MS, Bullimore J, Pretty JN, Osborn AM, Ball AS (2003) Soil type is the primary determinant of the composition of the total and active bacterial communities in arable soils. Appl Environ Microbiol 69:1800–1809. http://doi.org/10.1128/AEM.69.3.1800-1809.2003

Hashimoto S, Carvalhais N, Ito A, Migliavacca M, Nishina K, Reichstein M (2015) Global spatiotemporal distribution of soil respiration modeled using a global database. Biogeosciences 12:4121–4132. http://doi.org/10.5194/bgd-12-4331-2015

Halverson LJ, Jones TM, Firestone MK (2000) Release of intracellular solutes by four soil bacteria exposed to dilution stress. Soil Sci Soc Am J. 64:1630–1637.

He Y, Yang J, Zhuang Q, Harden JW, McGuire AD, Liu Y, Wang G, Gu L (2015) Incorporating microbial dormancy dynamics into soil decomposition models to improve quantification of soil carbon dynamics of northern temperate forests. J Geophys Res Biogeosci 120:2596–2611. https://doi.org/10.1002/2015JG003130

Hoeppner SS, Dukes JS (2012) Interactive responses of old-field plant growth and composition to warming and precipitation. Global Change Biol 18:1754–1768. https://doi.org/10.1111/j.1365-2486.2011.02626.x

Hurst CJ, Knudson GR, McInerney MJ, Stetzenbach LD, Walt MV (1997) Manual of Environmental Microbiology ASM Press. Washington, DC.

Illeris L, Michelsen A, Jonasson S (2003) Soil plus root respiration and microbial biomass following water, nitrogen, and phosphorus application at a high arctic semi desert. Biogeochemistry 65:15–29.

Joynt J, Bischoff M, Turco R, Konopka A, Nakatsu CH (2006) Microbial community analysis of soils contaminated with lead, chromium and petroleum hydrocarbons. Microbial Ecol 51:209–219. http://doi.org/10.1007/s00248-005-0205-0

Kaprelyants AS, Kell DB (1993) The use of 5-cyano-2, 3-ditolyl tetrazolium chloride and flow cytometry for the visualisation of respiratory activity in individual cells of Micrococcus luteus. J Microbiol Methods 17:115–122. https://doi.org/10.1016/0167-7012(93)90004-2

Koyama A, Steinweg JM, Haddix ML, Dukes JS, Wallenstein MD (2018) Soil bacterial community responses to altered precipitation and temperature regimes in an old field grassland are mediated by plants. FEMS Microbiol Ecol 94:fix156. https://doi.org/10.1093/femsec/fix156

Le Quéré C, Andrew RM, Canadell JG, Sitch S, Korsbakken JI, Peters GP,… and Keeling RF (2016) Global carbon budget 2016. Earth Syst Sci Data 8:605. http://doi.org/10.5194/essd-8-605-2016

Lee KH, Jose S (2003) Soil respiration and microbial biomass in a pecan—cotton alley cropping system in Southern USA. Agrofor Syst 58:45–54.

Lennon JT, Jones SE (2011) Microbial seed banks: the ecological and evolutionary implications of dormancy. Nat Rev Microbiol 9:119–130.

Lennon JT, Aanderud ZT, Lehmkuhl BK, Schoolmaster DR (2012) Mapping the niche space of soil microorganisms using taxonomy and traits. Ecology 93:1867–1879. https://doi.org/10.1890/11-1745.1

Liu W, Zhang Z, Wan S (2009) Predominant role of water in regulating soil and microbial respiration and their responses to climate change in a semiarid grassland. Global Change Biol 15:184–195. https://doi.org/10.1111/j.1365-2486.2008.01728.x

Manzoni S, Schaeffer S, Katul G, Porporato A, Schimel J (2014) A theoretical analysis of microbial eco-physiological and diffusion limitations to carbon cycling in drying soils. Soil Biol Biochem 73:69–83. https://doi.org/10.1016/j.soilbio.2014.02.008

Monson RK, Lipson DL, Burns SP, Turnipseed AA, Delany AC, Williams MW, Schmidt SK (2006) Winter forest soil respiration controlled by climate and microbial community composition. Nature 439:711–714. http://doi.org/10.1038/nature04555

Placella SA, Brodie EL, Firestone MK (2012) Rainfall-induced carbon dioxide pulses result from sequential resuscitation of phylogenetically clustered microbial groups. Proc Natl Acad Sci USA 109:10931–10936. https://doi.org/10.1073/pnas.1204306109

Salazar A, Sulman B, Dukes JS (2018) Microbial dormancy promotes microbial biomass and respiration across pulses of drying-wetting stress. Soil Biol Biochem 116:237–244. https://doi.org/10.1016/j.soilbio.2017.10.017

Salazar-Villegas A, Blagodatskaya E, Dukes JS (2016) Changes in the size of the active microbial pool explain short-term soil respiratory responses to temperature and moisture. Front Microbiol 7. http://doi.org/10.3389/fmicb.2016.00524

Schimel J, Balser TC, Wallenstein M (2007) Microbial stress-response physiology and its implications for ecosystem function. Ecology 88:1386–1394. https://doi.org/10.1890/06-0219

Schindlbacher A, Rodler A, Kuffner M, Kitzler B, Sessitsch A, Zechmeister-Boltenstern S (2011) Experimental warming effects on the microbial community of a temperate mountain forest soil. Soil Biol Biochem 43:1417–1425. https://doi.org/10.1016/j.soilbio.2011.03.005

Schindlbacher A, Wunderlich S, Borken W, Kitzler B, Zechmeister-Boltenstern S, Jandl R (2012) Soil respiration under climate change: prolonged summer drought offsets soil warming effects. Global Change Biol 18:2270–2279. https://doi.org/10.1111/j.1365-2486.2012.02696.x

Sinsabaugh RL, Turner BL, Talbot JM, Waring BG, Powers JS, Kuske CR, Moorhead DL, Follstad Shah JJ (2016) Stoichiometry of microbial carbon use efficiency in soils. Ecol Monograph 86:172–189. https://doi.org/10.1890/15-2110.1

Six J, Frey SD, Thiet RK, Batten KM (2006) Bacterial and fungal contributions to carbon sequestration in agroecosystems. Soil Sci Soc Am J 70:555–569. http://doi.org/10.2136/sssaj2004.0347

Stenström J, Svensson K, Johansson M (2001) Reversible transition between active and dormant microbial states in soil. FEMS Microbiol Ecol 36:93–104.

Stocker T (2014) Climate change 2013: the physical science basis: Working Group I contribution to the Fifth assessment report of the Intergovernmental Panel on Climate Change. Cambridge University Press, United Kingdom and New York, USA.

Suseela V, Conant RT, Wallenstein MD, Dukes JS (2012) Effects of soil moisture on the temperature sensitivity of heterotrophic respiration vary seasonally in an old-field climate change experiment. Global Change Biol 18:336–348. https://doi.org/10.1111/j.1365-2486.2011.02516.x

Waldrop MP, Firestone MK (2006) Seasonal dynamics of microbial community composition and function in oak canopy and open grassland soils. Microbial Ecol 52:470–479. http://doi.org/10.1007/s00248-006-9100-6

Wang WJ, Dalal RC, Moody PW, Smith CJ (2003) Relationships of soil respiration to microbial biomass, substrate availability and clay content. Soil Biol Biochem 35:273–284. https://doi.org/10.1016/S0038-0717(02)00274-2

Wang G, Mayes MA, Gu L, Schadt CW (2014) Representation of Dormant and Active Microbial Dynamics for Ecosystem Modeling. PloS one 9:e89252. https://doi.org/10.1371/journal.pone.0089252

Wang G, Jagadamma S, Mayes MA, Schadt CW, Steinweg JM, Gu L, Post WM (2015) Microbial dormancy improves development and experimental validation of ecosystem model. ISME J 9:1–12. http://doi.org/10.1038/ismej.2014.120

Waring BG, Hawkes CV (2015) Short-term precipitation exclusion alters microbial responses to soil moisture in a wet tropical forest. Microbial Ecol 69:843–854. http://doi.org/10.1007/s00248-014-0436-z

Wieder WR, Allison SD, Davidson EA, Georgiou K, Hararuk O, He Y, Hopkins F, Luo Y, Smith MJ, Sulman B, Todd-Brown K (2015) Explicitly representing soil microbial processes in Earth system models. Global Biogeochem Cy 29:1782–1800. https://doi.org/10.1002/2015GB005188

Zhang W, Parker KM, Luo Y, Wan S, Wallace LL, Hu S (2005) Soil microbial responses to experimental warming and clipping in a tallgrass prairie. Global Change Biol 11:266–277. https://doi.org/10.1111/j.1365-2486.2005.00902.x

Zhao Z, Peng C, Yang Q, Meng FR, Song X, Chen S, Epule TE, Li P, Zhu Q (2017) Model prediction of biome-specific global soil respiration from 1960 to 2012. Earth’s Future 5:715–729. https://doi.org/10.1002/2016EF000480

Zhou J, Xue K, Xie J, Deng Y, Wu L, Cheng X, Fei S, Deng S, He Z, Van Nostrand JD, Luo Y (2011) Microbial mediation of carbon-cycle feedbacks to climate warming. Nat Clim Change 2:106–110. http://doi.org/10.1038/nclimate1331

Zogg GP, Zak DR, Ringelberg DB, White DC, MacDonald NW, Pregitzer KS (1997) Compositional and functional shifts in microbial communities due to soil warming. Soil Sci Soc Am J 61:475–481.

